# Cellular-scale silicon probes for high-density, precisely-localized neurophysiology

**DOI:** 10.1101/2020.05.26.117572

**Authors:** Daniel Egert, Jeffrey R Pettibone, Stefan Lemke, Paras R. Patel, Ciara M. Caldwell, Dawen Cai, Karunesh Ganguly, Cynthia A. Chestek, Joshua D. Berke

## Abstract

Neural implants with large numbers of electrodes have become an important tool for examining brain functions. However, these devices typically displace a large intracranial volume compared to the neurons they record. This large size limits the density of implants, provokes tissue reactions that degrade chronic performance, and impedes the ability to accurately visualize recording sites within intact circuits. Here we report next-generation silicon-based neural probes at cellular-scale (5×10µm cross-section), with ultra-high-density packing (as little as 66µm between shanks) and 64 or 256 closely-spaced recording sites per probe. We show that these probes can be inserted into superficial or deep brain structures and maintain high-quality spike recordings in freely behaving rats for many weeks. Finally, we demonstrate a slice-in-place approach for the precise registration of recording sites relative to nearby neurons and anatomical features, including striatal µ-opioid receptor patches. This scalable technology provides a valuable tool for examining information processing within neural circuits, and potentially for human brain-machine interfaces.

## Introduction

Much of our current understanding of neural functions was gained through electrophysiological recording from individual neurons in behaving animals, one-at-a-time. Yet recording neurons one-at-a-time provides only a very limited view of information processing, which involves rapid interactions and coordination between neurons^12^. Simultaneous recordings from ensembles of many neurons has been achieved using large numbers of wires^1^, microfabricated electrode grids (e.g. the Utah array^2^), or microelectrode arrays shaped through photolithography^3–7^. These important approaches have yielded valuable results, but nonetheless share substantial limitations, especially for investigating densely-packed, locally-connected neurons.

To achieve the stiffness needed for a millimeters-long element to penetrate the brain, each element has typically had a width of at least 25-100µm (with cross-section in the thousands of µm^2^). This large foreign body causes substantial direct mechanical damage, and is also detected and rejected by the brain’s immune system^8–12^. The immune reaction leads to loss of neurons in the vicinity of the electrodes, one factor that frequently curtails the duration of chronic neural recordings^13^. The spacing between large elements must also be large to distribute tissue damage - for example, in a standard Utah array contacts are separated by 400µm. For many brain regions and cell types, this means that the neurons monitored from different electrodes are unlikely to be in direct communication with each other^14,15^.

Compounding this problem, large devices must be removed from the brain before processing the tissue for histological analysis. Since the recording electrodes are not observed *in situ*, it has proven difficult to obtain accurate registration between electrophysiological recordings and key anatomical markers. Lack of anatomical registration has impeded research in several subfields, including the investigation of the function of striatal “patches” (irregularly-shaped, ∼100µm-scale zones that are detectable through staining for mu-opioid receptors; aka striosomes^16,17^). Although visualization of spatial patterns of ensemble activity can be achieved through other methods - such as multiphoton imaging of genetically-encoded calcium indicators – these currently lack the exquisite temporal resolution of electrophysiology (though see^32^).

Several approaches have been used to insert smaller electrodes into the brain. For example, flexible, cellular-scale electrodes can be infused through a micropipette^18,19^ and linear polymer probes can be dragged into the brain using a stiff needle or silicon support as a shuttle^20,21^. However these insertion devices are typically at least as large as traditional electrodes (25µm+ in width; though see^22^. The resulting acute “stab-wound” has been shown to cause substantial, permanent neuron loss even if no foreign body remains in the brain^12^. As an alternative, very thin (<10µm) electrodes made from carbon fibers can be directly inserted, either one at a time^18^, in bundles^24^ or mounted on silicon supports to increase overall stiffness and penetration^23^. The latter approach shows promising chronic performance^13^, but has not yet demonstrated the feasibility of large channel counts.

To overcome these limitations, we designed neural probes with high channel counts but considerably smaller physical shank dimensions than standard silicon devices. We reasoned that the force required to insert an element without buckling depends on its length; by coupling a relatively short (500µm) electrode-bearing lower portion with cellular-scale cross-section (5×10µm) to a more robust upper portion (far from the recording sites), we obtained reliable insertion even deep into the brain. The smaller size probes allow much higher density neural recording, with excellent chronic performance for months (at least) after implant. We further developed a method whereby probes are left in place during brain sectioning and histological processing, by slicing through the entire head after decalcifying the skull. This was made possible by the very thin probe shanks that can be easily sliced through without distorting the surrounding tissue. By cutting thick (∼300µm+) slabs of tissue and using immunohistochemistry to detect key anatomical features, we found that we can directly visualize identified recording sites within intact micro- and meso-scale neural circuits.

## Results

### Neural probe design

Each device consists of 32 shanks in a comb-like configuration (Fig. 1). Each shank has an electrode-bearing tip section (500µm long) with cellular-scale cross-section (10µm wide x 5µm thin). We designed multiple probe variations, in which each shank has lengths of 3-9mm, and holds two or eight electrode sites (64 or 256 total channels / probe). Adjacent shanks differ in length by 50µm to reduce peak insertion force. To enable insertion, probes have a wider upper portion, with 25µm width and 11.5µm thickness. For 6mm and 9mm long shanks, this upper portion widens gradually to 40µm at the top.

**Figure 1:**
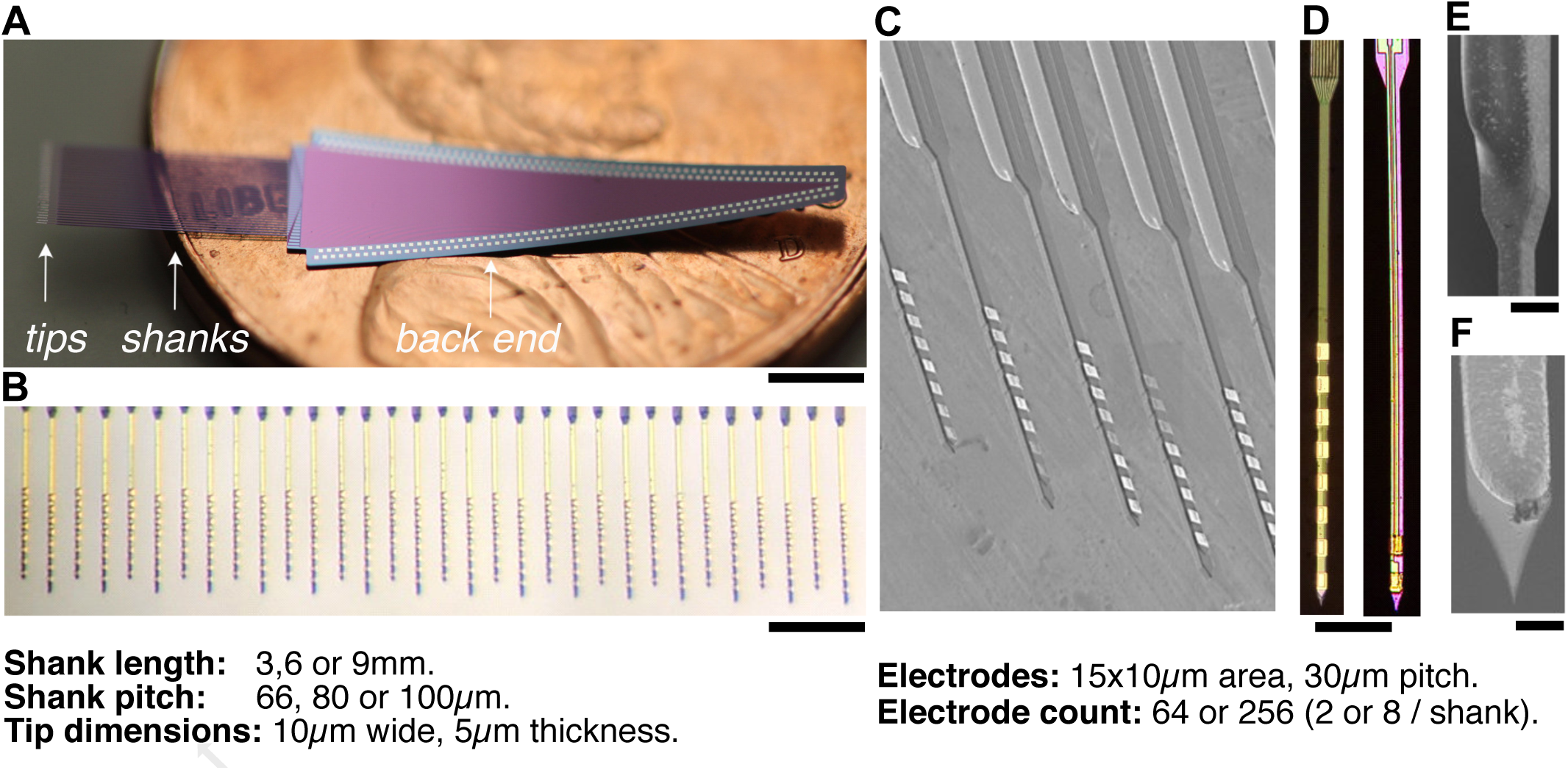
Silicon probes with cellular-scale tips. **A**, 256-channel probe (6mm shank version), on penny for scale. Scale: 3mm. **B**, Close-up of tip portion showing 32 shanks, each with 8 electrodes. Scale: 200*µ*m. **C**, Electron micrograph of probe tips, showing electrodes. **D**, Comparison of shanks with 8 electrodes (left; in 256 channel probes) and 2 electrodes (right; in 64 channel probes). Scale: 50*µ*m. **E**, Smooth transition to stiffener-reinforced upper section. Scale: 10*µ*m. **F**, Sharpened tip. Scale: 5*µ*m.

### Nanofabrication Process

We used a bulk silicon fabrication process to shape the shanks, and thin film technology for interconnects, electrodes and insulators (Fig. 2). Materials and processing steps were derived from the Michigan probes process^25^ which uses diffusion of boron into silicon to form etch stops and pattern the probe shape. Each probe design used one or more of the following technical refinements:

**Figure 2:**
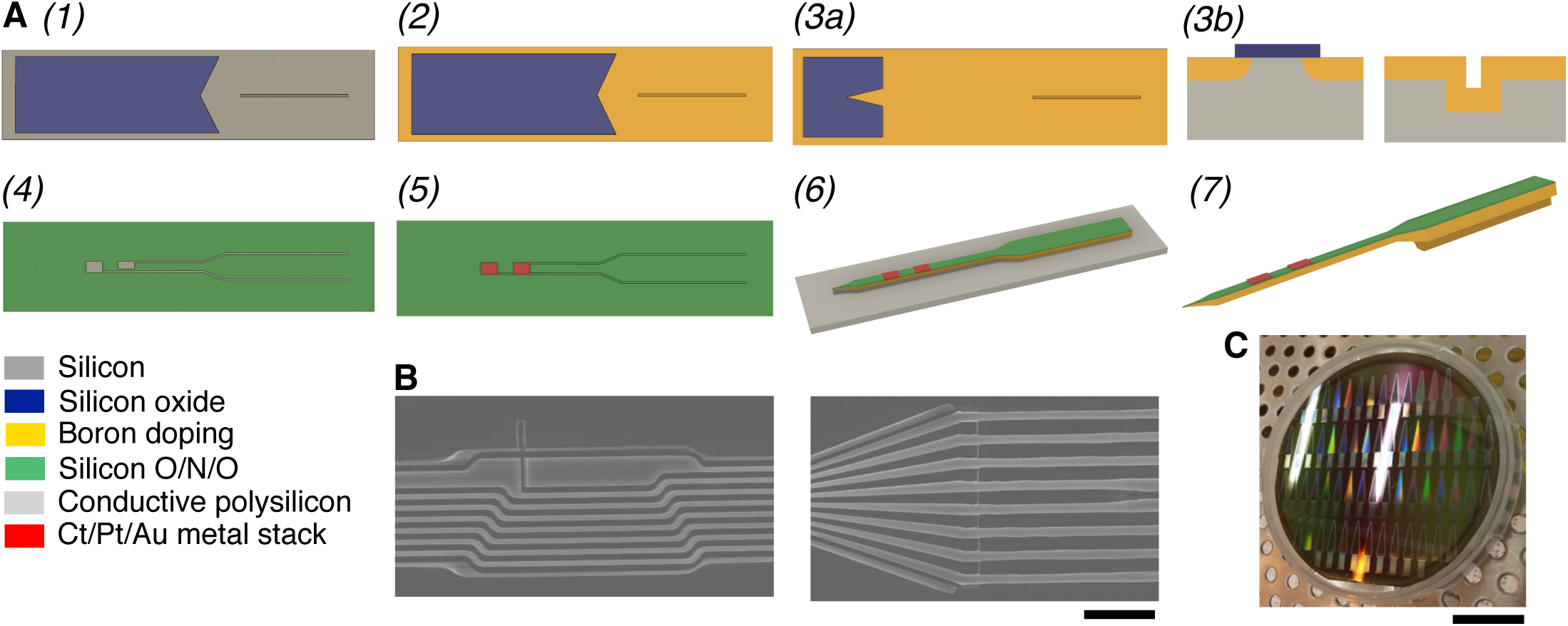
**A**, Main steps and materials of the nanofabrication process flow: *(1)* Grow and pattern silicon oxide to form a masking layer. Etch trench using deep reactive ion etching (DRIE) for stiffeners. *(2)* Diffuse boron into silicon in places that were not masked. *(3)* Remove the mask except for a small section around the tip ends. Perform a second boron diffusion to define tip thickness *(3a top view, 3b side view). (4)* Deposit stack of electrically insulating silicon oxide, silicon nitride and silicon oxide. Deposit layer of electrically conductive polysilicon and pattern to form interconnects. *(5)* Deposit another insulator stack (as before). Etch vias, sputter Cr/Pt/Au and pattern via lift-off to form bonding pads and electrodes. *(6)* Etch probe outline using DRIE. *(7)* Etch undoped silicon with ethylene diamine pyrocatechol (EDP). **B**, Close-up of polysilicon interconnects shaped with e-beam lithography (left) and transition to stepper patterning (right). Scale: 5*µ*m. **C**, Completed wafer. For handling and protection of the fine tips the devices remain attached to the wafer frame by small tabs. Scale: 2cm.

First, we patterned very high-density traces within the tip sections via electron-beam lithography, which allows for orders-of-magnitude smaller features^26^ compared to optical patterning. As e-beam lithography is a serial and thus relatively expensive process, we used it only on the lower, thin tip sections of 256-channel devices, to define the narrow valleys separating traces. The remaining traces were patterned using conventional optical stepper lithography.

Second, we integrated a stiffener into upper shank sections by selectively increasing thickness there (6mm, 9mm shank versions only). Traditionally in this process, neural probes have a uniform thickness defined by the depth of diffused boron in silicon. Here, a trench was etched along parts of the shank before boron diffusion. The sidewalls and the bottom of the trenches were exposed during boron doping and hence the thickness of this part of the shank is extended by the depth of the trench (increasing maximum thickness to 30µm). The trenches were kept sufficiently narrow such that they were completely refilled in subsequent steps.

Third, we produced sharper tips by masking boron diffusion towards the end of the shank (3mm shank version only). Increasing tip sharpness can reduce compression of the brain during insertion and make buckling of the shanks less likely^27^. By masking boron diffusion at the very tip, the thickness of the released probe was reduced to that of the remaining insulating layers, around only 1.5µm.

### Successful implantation into shallow and deep brain structures

After wire-bonding to custom PCBs, each finalized probe design was found to successfully insert into brain without requiring specialized surgical practices. During the design stage of this project, test probes were fabricated with 5mm length, without stiffeners and with a uniform width of the upper section. These were found to have an insertion yield of 75% (42/56 shanks). Adding stiffeners and the tapering increase in top section width improved this yield to 97% (36/37 shanks).

Fig. 3A shows images of inserted devices with 3mm long shanks, with an extremely narrow pitch of 66µm during implantation 1.5mm deep into rat cortex. We were also able to insert multiple devices, mounted on thin flexible polyimide cables, into the same hemisphere, targeting both motor cortex with a 6mm probe and striatum with a 9mm probe (Fig. 3B). Once inserted, the probes were secured, the site was sealed (see Methods), and rats consistently recovered from surgery (Fig. 3C) without behavioral impairments.

**Figure 3:**
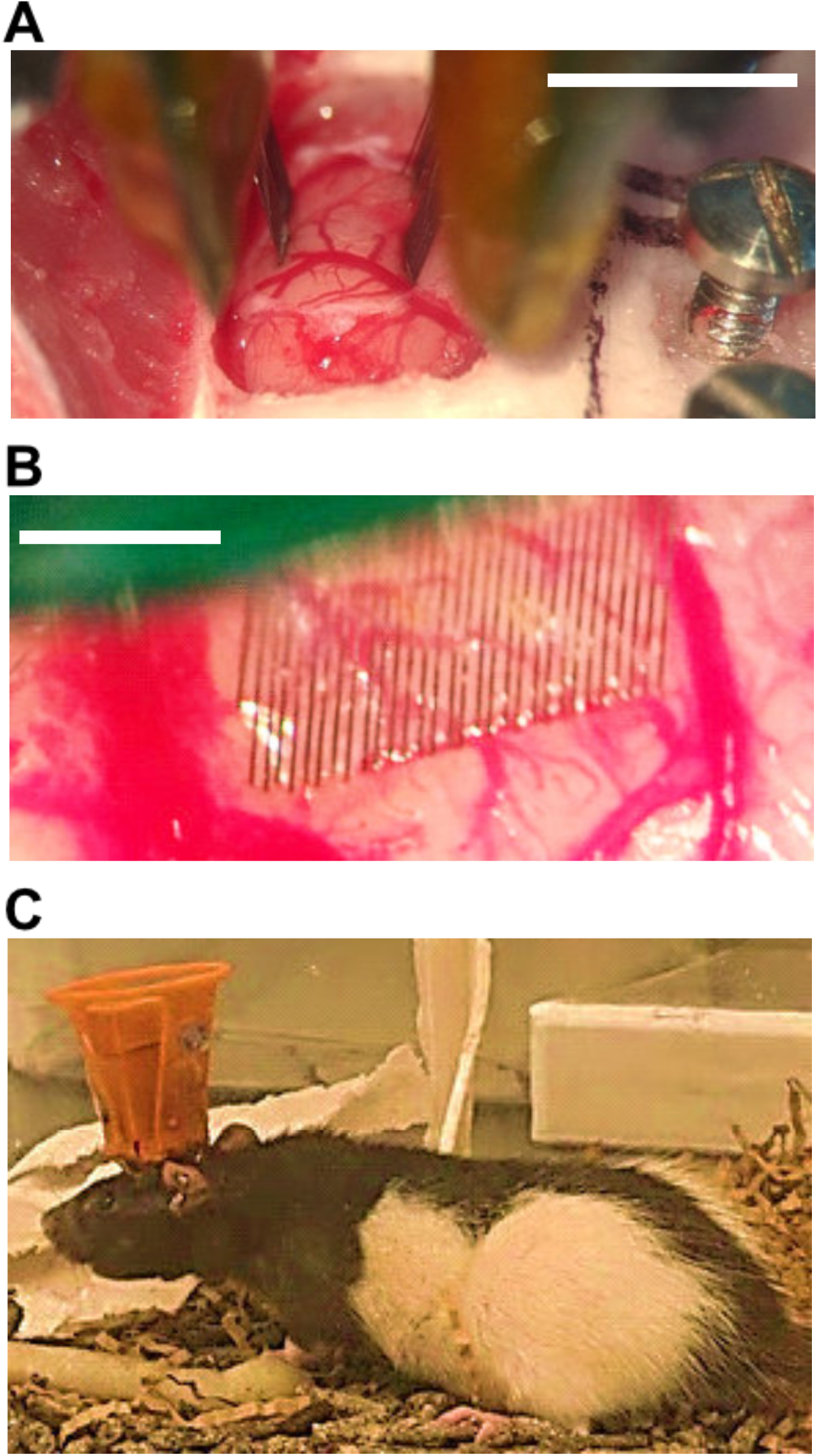
Probe insertion. **A**, Two devices (6mm, 9mm lengths) being implanted into the same hemisphere for simultaneous recording from motor cortex and striatum. Scale: 2mm. **B**, Device with 66*µ*m pitch between shanks (and 3mm length) being inserted 1.5mm deep into motor cortex. Scale: 1mm. **C**, Rat after recovery, with 256 channel implanted device connected to 2 x 128 channel lntan headstages within a 3D-printed enclosure.

### Recording from large neuronal populations

We successfully implanted 256-channel devices into brain structures both superficial (motor cortex) and deep (striatum, globus pallidus (GP)), and obtained high-quality chronic spike recordings in freely-moving rats from each structure. Fig. 4A shows an example from a 9mm, 256-channel probe targeting GP (3 days post-surgery), yielding 259 distinct simultaneously-recorded single-unit clusters.

**Figure 4.**
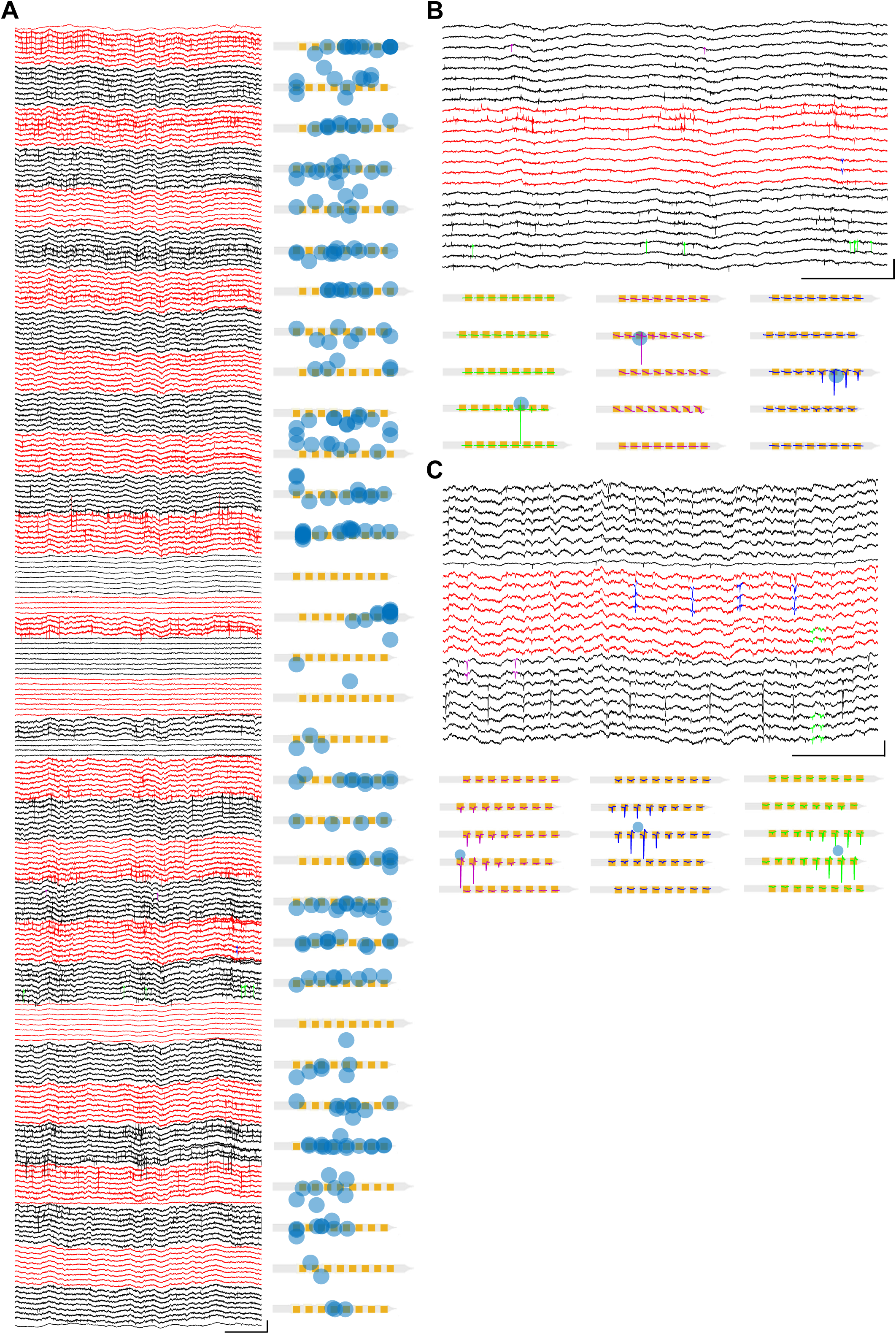
High-density chronic spike recording. Left, example of signals from a 256-channel probe (100*µ*m shank pitch) targeting rat GP (depth 6.5mm), 3 days post-surgery. Traces are 250ms duration, unfiltered and grouped by shank (red / black). Right, spike sorting for this recording session produced 259 clusters. Each cluster is marked by a blue circle at its estimated physical location (see Methods). Scale bars: 1mV / 50ms. **B**, Illustration of the waveform distribution across electrodes for three example neurons (purple, blue, green) from the same session as A. Top shows waveforms highlighted in wide-band signals (250ms duration); bottom shows corresponding average waveforms, each visible only on a small subset of electrodes. The top traces are from the middle three shanks in the bottom illustration. Scale bars: 1mV / 50ms. **C**, As B, but recording from a 256-channel probe (66*µ*m shank pitch) placed in rat M1 cortex (18 days post-surgery). M1 cells typically generated wider electric fields, visible across many electrodes over two to three adjacent shanks. Scale bars: 1mV / 50ms.

The dense electrode arrangement both within- and between-shanks allows examination of the spatial extent of extracellular voltage changes associated with action potentials from each neuron. In our GP example (using 100µm shank pitch), each averaged action potential was visible only across a small number of recording sites (Fig. 4B). This is consistent with relatively small extracellular fields generated by these neurons, which typically have radially-symmetric dendritic fields^28^. By contrast, when we recorded from motor cortex (using a probe with just 66µm shank pitch), each averaged action potential was readily visible across many sites. This includes both sites along a shank (consistent with recording along the large apical dendrite^29^) and across two or even three shanks (Fig. 4C). This suggests that – at least for recording cortical projection neurons - our probes have sufficient density to detect spikes from a large proportion of cells within the two-dimensional field of recording sites.

### High-density, chronic spike recordings

We were able to obtain high-quality spike recordings from cellular-scale silicon probes over extended time periods: Fig. 5A shows an example recording at 139 days post-implant. To quantify changes in chronic performance over time we recorded from a cohort of rats (n=4) each with a 64-channel probe implanted in dorsal striatum, at 1 week and 7 weeks post-implant (Fig. 5B).

**Figure 5:**
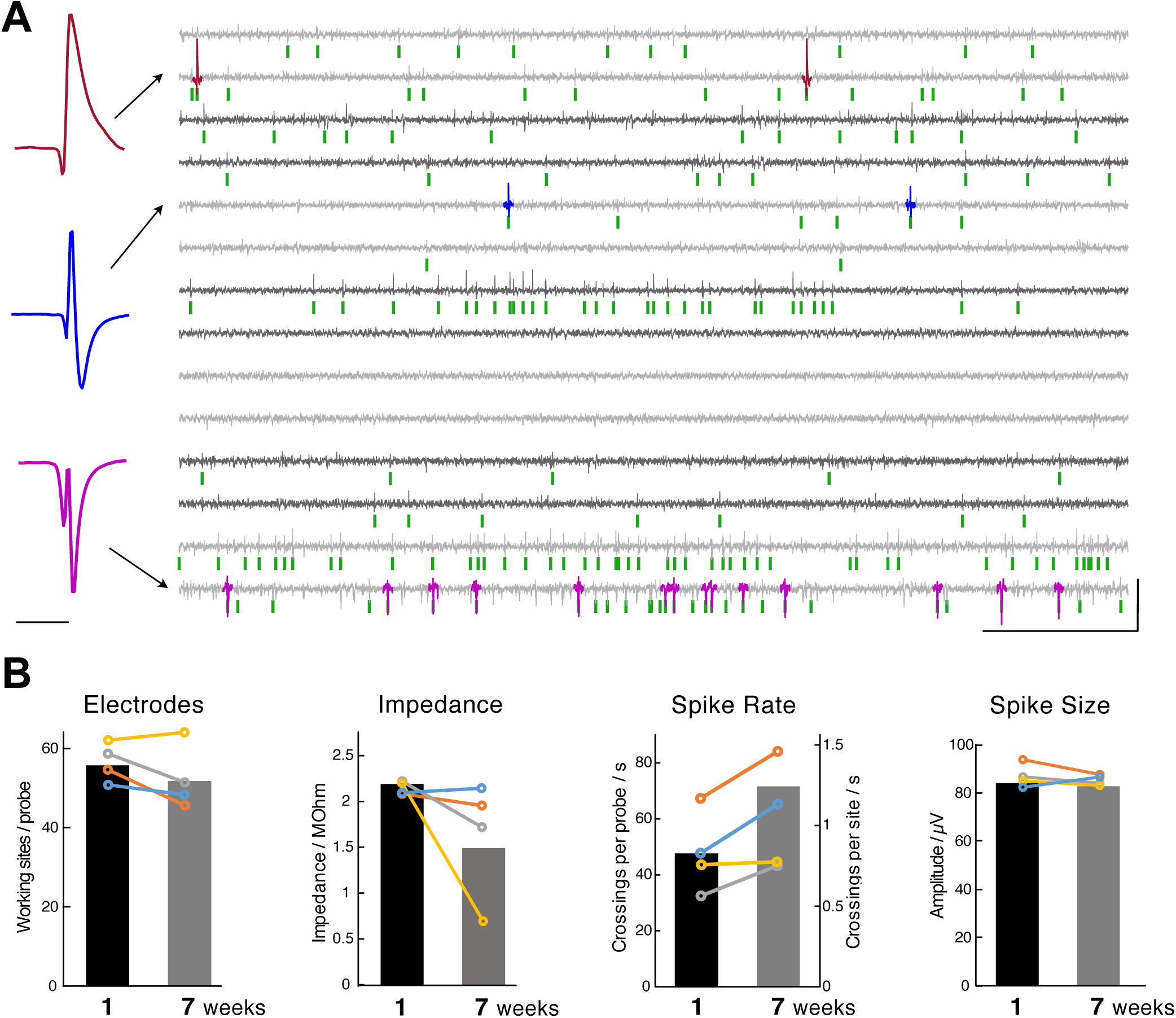
Cellular-scale silicon probes record spikes for many weeks. **A**, Example recording from a 64-channel probe, 139 days after implantation into dorsal striatum (250ms shown). Signals are high-pass filtered and color-coded by shank (alternating light and dark gray; 7 shanks shown out of 32 total). Examples of spikes from large units are highlighted in color, with the corresponding average waveforms shown at left (peak-valley ranges: brown 300*µ*V, blue 186*µ*V, magenta 190*µ*V). Green marks below signals indicate all detected spike events (threshold of 6 x median absolute deviation). Scales (left): 1ms; (right) 300*µ*V / 50ms. **B**, Performance metrics comparing 1 week post-implant (6-7 days) with 7 weeks (49-50 days). Same implant type, same 6 x M.A.D. event threshold as in (A). Colored lines indicate individual animals (n=4 probes, 256 sites total), black and grey bars indicate probe means (in left-most plot) or site means (other plots).

We saw a slight decline (from 55.8 to 52.0, out of 64) in the mean number of working electrodes (defined as those with impedances between 0.5 and 4.0MOhm); the remaining sites showed reduced impedance (mean per device fell from 2.2MOhm to 1.6MOhm; mean per site from 2.2MOhm (n=225) to 1.5MOhm (n=211); two-tailed Wilcoxon signed-rank test, p=1.3e-19). The mean rate of spike events increased (per device: from 47Hz to 60Hz threshold crossings/s/device; per site: from 0.8Hz (n=225) to 1.2Hz/s (n=211); two-tailed Wilcoxon signed-rank test for sites recorded at both timepoints, p=0.0481). The size of detected spikes was stable (mean per device, 87.0µV to 85.4µV; mean per site 84.0µV (n=225) to 83.0µV (n=211); no change in size for those recorded at both timepoints; two-tailed Wilcoxon signed-rank test, p=0.2425). We conclude that these devices are able to consistently record spikes over many weeks.

### Anatomical registration of implanted probes *in situ*

To facilitate registration between recording sites and microcircuit features, we wished to avoid pulling electrodes out of the brain at the end of experiments, and instead leave electrodes *in situ* during processing for histology. Many standard electrodes (e.g. tetrodes) cannot be cut through by cryostat blades without extensive tissue damage (J.R.P. and J.D.B, unpublished observations). Some recent electrode designs can be cut through^30^ but do not preserve the orderly arrangement of electrodes needed for registration between specific recording channels and histological locations.

We found that – following formaldehyde perfusion and decalcification of the skull – we can cleanly cut through skull, brain, and probe shanks of rats with implanted cellular-scale silicon probes (Fig. 6A,B). By cutting 300µm slabs we could directly observe the electrode-bearing portion of the shank tips within brain tissue (Fig. 6B), while still allowing antibodies to fully penetrate the tissue for immunohistochemistry (Fig. 6C). Staining for the neuronal marker NeuN revealed many neuronal cell bodies in close proximity to the tips of chronically-implanted probes (Fig. 6D), consistent with minimal damage to local circuits. Staining for µ-opioid receptors (MOR) revealed striatal patches (Fig. 6C,D), and we were able to use this marker to perform 3-D reconstructions of shank tips *in situ* within patches (Fig. 6E). This slice-in-place approach thus allows effective direct visualization of electrodes within intact brain circuits.

**Figure 6:**
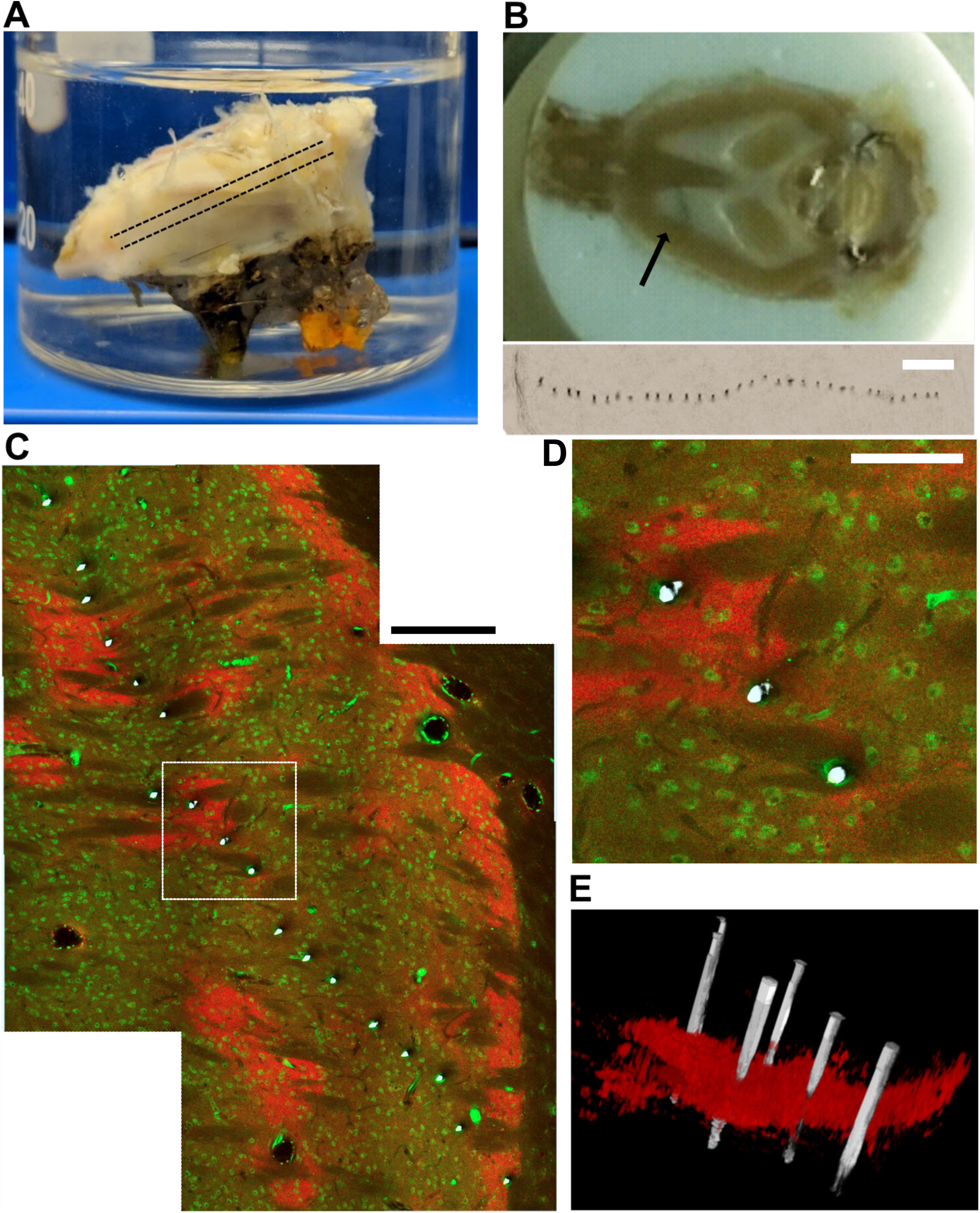
Slice-in-place approach for precise localization of electrodes within brain circuits. **A**, Decalcification of the skull (shown upside-down) after removal of superficial tissue. The dashed line indicates the plane and thickness of slices. **B**, Top, skull and brain are sliced together, ventral to dorsal, with the probes *in situ*. Faint gray line (arrow) is the row of shanks. Bottom, example of probe tips within striatum, at about 4.3 mm depth. Scale: 400*µ*m. **C**, Brightfield-imaged shank tips (white) superimposed on immunofluorescence imaging of neurons (NeuN stain, green) and striatal patches (MOR; *µ*-opioid receptors, red) in the same plane. Scale: 200*µ*m. **D**, Close-up of rectangle area in C. Scale: 100*µ*m. **E**, 3D visualization of shank tips (white) relative to a MOR-stained patch.

## Discussion

We have demonstrated that silicon probes with cellular-scale tip sections can yield long-lasting chronic spike recordings in freely-moving animals, large numbers of densely-recorded neurons, and direct visualization of probe tips relative to histological markers. This combination of features is likely to be very useful for many systems neuroscience studies, especially investigations of neural circuit functions over a range of spatial scales.

The reduced damage and improved localization possible with very thin shanks are advantages compared to alternative advanced silicon probe designs, which have typically emphasized large numbers of sites on wider shanks (e.g. ^6,29,31,32^). Those designs may nonetheless have advantages for some applications, such as when multiple targets are arranged vertically within the brain^6^. In other situations, such as recording from a horizontal cell layer, the current design may be more appropriate.

Neural probes made using flexible polymer substrates^20,22,30,33^ can also produce recordings with high longevity. Furthermore, their mechanical flexibility prevents breaking, and they may also be sliced-in-place^34^. However, their insertion often requires transient mechanical support within the brain by removable shuttles^35,36^, that cause acute (stab-wound) damage and hinder narrow spacing between shanks. Once implanted, flexible devices cannot be further moved^37^, preventing adjustments based on observed electrophysiological features after recovery, or recording of new sets of cells. Further advantages of silicon include the sophisticated assembly tools developed for silicon technology, and the potential to seamlessly include active electronics, for example for multiplexed readout^6^, or LEDs for optical stimulation^38^.

An alternative technology for recording many neurons simultaneously is optical imaging. This can have several important advantages including direct observation of neuron locations, visualization of distinct neuronal compartments (such as dendrites^39^), or wide fields-of-view^40^. However, imaging of large numbers of neurons has to date typically examined superficial structures in head-fixed animals, with calcium fluctuations detected by GCaMP as the activity measure. Calcium dynamics can have a non-straightforward relationship to spiking, and action potentials remain the gold standard measure of neural “activity”. Imaging action potentials is possible using voltage sensors instead^39^ but the light intensity requirements for imaging at near-millisecond resolution make large fields of view impractical with current sensors. Recording from deep brain structures in freely moving animals has been made possible using miniaturized microscopes and gradient refractive index lenses^41^. However these lenses are typically 0.5-2mm in diameter, and implantation involves aspirating overlying brain structures, potentially altering neural dynamics.

We deliberately placed recording electrodes far from the relatively thicker, stiffer shank portions, which presumably provoke greater tissue reactions. How far away is far enough? For solid silicon probes with 300µm width and 15µm thickness, Skousen *et al*.^42^ showed that after 2 months neural density is low in the immediately adjacent vicinity, but is normal at 100µm distance. In the present design the closest recording site is 250µm away from the thicker section (for 256-channel devices; 430µm away for 64-channel devices). We saw no indication that the electrodes further away from the tip (and thus closer to the thicker sections) record fewer neurons (Fig. 5A), suggesting that we have achieved sufficient distance from any damage caused by thicker upper sections. Although the aggregate cross-section of our recording tip sections is substantial (1600µm^2^, comparable to the single shank of Neuropixels probes), distributing this displacement over many small elements has been shown^42^ to avoid deleterious tissue reactions.

An important objective was to improve co-registration of recording sites with meso-scale histological markers. We and others have previously attempted co-registration with µ-opioid-receptor-defined striatal patches by observing tissue damage or gliosis produced by electrodes^38,43^, passing current to produce marker lesions^44,45^ or coating probes in lipophilic tracer dyes such as DiI^46^. We have not found these approaches to be highly reliable and effective for identifying patch neurons – for example, dye coating is typically too uneven, while lesions destroy the essential immunochemical markers just where they are most needed. Some studies of patch neurons identified using imaging have begun to appear^47,48^. However, our slice-in-place approach offers higher temporal resolution with less damage to nearby brain tissue.

We chose 300µm thick slabs for slice-in-place analysis. Thinner sections would include less of the recording zone and require more registration between sections, and increase the chance that short shank portions fall out of tissue during slicing. Thicker slabs would avoid these concerns, but are harder for antibodies to penetrate, and make microscopy more challenging due to increased light scattering. Scattering can be greatly reduced by tissue clearing^35,49,50^ but we have found it is hard to avoid measurable shrinkage/expansion as the tissue is cleared, which impedes co-registration. Future improvements to our approach could use thicker slabs (or whole brains) together with enhanced clearing methods that avoid changes in tissue volume, and smaller antibodies (e.g. nanobodies^51^) for more effective tissue penetration. The small size of the tips of the shanks might lend itself well to forming three-dimensional electrode arrangements, for example by stacking^32^. If brain structures at different depths are targeted for simultaneous recording, the length of each shank and arrangement of can be chosen individually, and clusters of shanks can be precisely tailored to their targets.

Overall the cellular-scale silicon probes presented here are part of a new generation of devices for chronic monitoring of large neural populations^6,20,21,52^. Their complementary geometry, small feature size, and the ability to achieve co-registration with histology make these probes suitable for addressing many outstanding questions in systems neuroscience.

## Methods and Materials

### Nanofabrication process flow and probe assembly

The probes were fabricated at the Lurie Nanofabrication Facility at the University of Michigan. The process begins with thermally growing and patterning a 1.2µm thick oxide on a silicon wafer as a masking layer. A 20µm deep and 1.2µm wide trench is etched using DRIE (deep reactive ion etching) where stiffeners are placed. The etch recipe was optimized to achieve vertical side-walls and high aspect ratio trenches. Boron is thermally diffused at 1175°C, 12µm deep into silicon surface in places that were not masked. The mask is removed except for a small section around the probe tip ends. A second, 5µm shallow boron diffusion is performed to define the thickness of the tips. A stack of electrically insulating silicon oxide and silicon nitride is deposited using low pressure chemical vapor deposition. A 1.4µm thick layer of electrically conductive, phosphorous-doped polysilicon is deposited and patterned to form interconnects. For this step, patterns formed by stepper and e-beam lithography (for 256-channel probes) are stitched together. The e-beam was only used to keep the narrow traces separate. The polysilicon traces routing along the tip of the probes were designed to have a 470nm minimum feature size. Another stack of insulators (identical to those previously mentioned) is deposited. After etching vias, Cr/Pt/Au is sputtered and patterned via lift-off to form bonding pads and electrodes. Lastly, the outline of the probe is defined using DRIE, and any undoped silicon is wet etched in EDP (ethylenediamine pyrocatechol). For handling and protection of the fine tips, the devices remain attached to the wafer frame by small tabs. The probes were mounted onto printed circuit boards (PCBs) using Crystalbond 509 (SPI supplies) and wire bonded.

### Animal Surgery

All animal procedures were approved by the Institutional Animal Care and Use Committees at the University of Michigan and the University of California, San Francisco. The probes were implanted into adult male Long-Evans rats, weighing 300 – 350g. Anesthesia was initialized with 5% isoflurane (v/v). The rats were maintained under isoflurane anesthesia, which was continuously monitored using toe pinch and breathing rate, and the flow of isoflurane was adjusted accordingly. The head was shaved at and around the area of the incision site. The shaved area was swabbed using alternating applications of betadine and 70% ethanol. Ointment was applied to the eyes to keep them from drying during surgery. Ear bars were mounted in both ears and fixed in a stereotaxic frame (Kopf Instruments, Model 900). After making an incision, the skin flaps were pulled apart using hemostats and the skull surface was cleaned using cotton swabs and 2% hydrogen peroxide (v/v). A burr bit (19008-07, Fine Science Tools, Foster City, CA) was used to drill holes around the periphery of the skull for bone screws (19010-00, Fine Science Tools, Foster City, CA). Reference and ground wires originating from the implant were attached to bone screws using MillMax pins and placed 1mm caudal to lambda and over the contralateral cerebellum, respectively. Next, a 2×4mm craniotomy was made over the target brain region. The dura was gently resected using dural forceps and hook, and insertion was performed within minutes of resection to avoid excessive swelling. Special care was taken to avoid larger blood vessels on the surface of the brain and if necessary, specific shanks were broken off. Just before the probe tips contacted the surface of the brain, excess liquid was removed in order to prevent shanks from wicking together. Shanks were lowered at about 100 µm/s into the brain. Striatal shanks were lowered to ∼4.2 mm below the surface of the brain, leaving ∼800 µm shank length above the brain surface. Motor cortical shanks were lowered ∼1.5mm below the surface (at +0.5AP, 3.5ML relative to bregma). For the cohort of rats used for chronic stability testing, and the pallidal implant, the exposed surface of the brain was sealed with Kwik-Sil (World Precision Instruments, Sarasota, FL)^23^. The motor cortex implant was sealed with cyanoacrylate glue (Loctite SuperGlue Gel control). The rat implanted in striatum for slice-in-place was sealed with a thin layer of DOWSIL (3-4680, Dow Corning, Midland, MI) and petroleum jelly (Vaseline) coated along the shanks^6^. Finally, the skull was covered with dental acrylic (Hygenic Acrylics, Switzerland).

### Recordings, impedance measurements

PCBs with 64 channel probes were connected to RHD2216 64-channel headstages, and PCBs with 256-channel probes were connected to two RHD2000 128-channel headstages (Intan Technologies, Los Angeles, USA). Recordings were made using the Intan interface software (version 1.5.2). The sampling rate was 30 kS/s for the striatal implants, 25 kS/s for the GP implant and 20 kS/s for the motor cortex implant. The sample depth was always 16-bit. Semi-automated clustering was performed using KiloSort2^53^ followed by manual curation using the ‘phy’ gui (https://github.com/kwikteam/phy). Impedances (at 1kHz) were measured using the Intan headstages. For assessment of spiking across repeated sessions, we examined brief (10 min) epochs as rats were resting quietly. Signals were first subjected to common average referencing, then high-pass filtered at 234Hz using a wavelet transform^54^. In line with prior studies^6^, spike events were detected using a threshold of 6x median absolute deviation of the signal, with 0.5ms dead time between possible spikes. To prevent contribution from occasional large movement artifacts, portions of the filtered signal that crossed a 1mV threshold were discarded, as were spike events that fell within a millisecond window in which the mean amplitude across all channels was more than 3x the average for the whole session. Analyses did not include channels that were not connected to brain (impedance outside the range 500-4000 kOhm) due to mechanical failure (e.g. detachment of connector pins after multiple plug-in, plug-out cycles). We estimated the physical location of recorded single-units in each of two dimensions, by finding the peaks of Gaussians fit to average peak spike amplitudes across the electrodes (averaged along, or across, shanks, respectively).

### Slice-in-place, histology and imaging

Rats were transcardially perfused with 4% paraformaldehyde (PFA) in phosphate buffered saline (PBS). After perfusion, the rats were decapitated, the skull was stripped of gross tissue, and the jaw was removed. The remaining bone, brain and probes were immersed in a solution of 0.25M tetra-sodium ethylenediaminetetraacetic acid (EDTA) in PBS, pH-balanced to 7.4. The solution was exchanged daily, with the soak continuing for 2-3 weeks until the bone turned rubbery. The skulls were soaked in 30% sucrose in PBS for 72hrs for cryoprotection, then the solution was progressively exchanged with OCT (optimal cutting temperature compound). Once the sample was immersed in 100% OCT, it was brought into a vacuum to facilitate penetration into ventricles then frozen using dry ice. Slicing was performed on a cryostat at -16°C. The skull was mounted such that the most ventral portion was accessed first. The tissue was sliced together with the bone and the probes, at 300µm thickness. The resulting slabs were washed, blocked and incubated for 7-10 days at room temperature with both primary antibodies Rb ∝ mOR (ImmunoStar 24216) and Ms ∝ NeuN (Millipore MAB377). The slabs were then incubated with secondary antibodies for 3-5 days at 4°C. Before imaging on a confocal microscope (Nikon TI), the slabs were soaked in a refractive index matching agent (TDE)^55^. The obtained images were processed using ImageJ and the plugin “3D Viewer”^56^ was used to generate the volume view, with a brightness-based threshold defining patch boundaries.

## Author Contributions

Conceptualization: DE, JDB; Methodology: DE, JRP; Formal Analysis: DE; Investigation: DE, SL, PRP, CMC, JRP; Writing – Original Draft: DE, JDB; Supervision: JDB, CAC, KG; Funding Acquisition: JDB, CAC, KG.

## Acknowledgements

Probe fabrication was performed at the Lurie Nanofabrication Facility, a member of the National Nanotechnology Infrastructure Network, which is supported in part by the National Science Foundation. This work was supported by NIH BRAIN Initiative grants U01NS094375 and UF1NS107659, the University of Michigan, and the University of California San Francisco. We thank Michael Farries for assistance with GP recordings.

